# midiaPASEF maximizes information content in data-independent acquisition proteomics

**DOI:** 10.1101/2023.01.30.526204

**Authors:** Ute Distler, Mateusz Krzysztof Łącki, Michał Piotr Startek, David Teschner, Sven Brehmer, Jens Decker, Thilo Schild, Jonathan Krieger, Florian Krohs, Oliver Raether, Andreas Hildebrandt, Stefan Tenzer

## Abstract

Data-independent acquisition (DIA) approaches provide comprehensive records of all detectable precursor and fragment ions. Here we introduce midiaPASEF, a novel DIA scan mode using mobility-specific micro-encoding of overlapping quadrupole windows to optimally cover the ion population in the ion mobility-mass to charge plane. Using overlapping ion mobility-encoded quadrupole windows, midiaPASEF maximizes information content in DIA acquisitions which enables the determination of the precursor m/z of each fragment ion with a precision of less than 2 Th. The Snakemake-based MIDIAID pipeline integrates algorithms for multidimensional peak detection and for machine-learning-based classification of precursor-fragment relationships. The MIDIAID pipeline enables fully automated processing and multidimensional deconvolution of midia-PASEF files and exports highly specific DDA-like MSMS spectra which are suitable for *de novo* sequencing and can be searched directly with established tools including PEAKS, FragPipe and Mascot. midiaPASEF acquisition identifies over 40 unique peptides per second and provides powerful library-free DIA analyses including phosphopeptidome and immunopeptidome samples.

## 1 Introduction

Since their introduction in the early 2000s, data-independent acquisition (DIA) methods have rapidly evolved and undergone major developments. Recent advancements in instrumentation, data acquisition schemes as well as novel algorithms for data analysis have greatly expanded the sensitivity and achievable proteomic depth and throughput of DIA-based workflows and address the requirements for reproducible explorative measurements in large sample cohorts [1, 2]. In contrast to data-dependent acquisition (DDA) schemes, where distinct precursor ions are selected according to their abundances and fragmented in a serial manner, DIA approaches record fragment ion information for each precursor in an unbiased way. As fragment ion information is acquired for all precursor ions within predefined mass-to-charge (m/z) windows regardless of their intensity or other characteristics, DIA overcomes inherent limitations of DDA-based workflows such as stochastic and irreproducible precursor ion selection and under-sampling [3]. The reduction of missing values in large sample cohorts and high-throughput studies markedly increased reproducibility and data completeness in DIA-based studies [1, 4]. However, DIA approaches face a different challenge as - in contrast to DDA - fragment ions are typically acquired using larger quadrupole windows, thus reducing the specificity of association between a fragment ion and its precursor. Therefore, DIA generates highly multiplexed fragment ion spectra, which require deconvolution and the high spectral complexity of DIA data still poses a significant challenge for downstream analysis. One option to reduce the spectral complexity is the integration of ion-mobility separation (IMS) into liquid chromatography-mass spectrometry (LC-MS)-based workflows. IMS provides an additional dimension of separation and increases overall system peak capacity, thereby improving both sensitivity and precursor ion selectivity while concomitantly reducing chimeric and composite interferences [3, 5, 6, 7].

In bottom-up proteomics, peptides are typically separated by reverse-phase LC with chromatographic elution peak widths in the range of seconds while spectral acquisition times on time-of-flight (TOF) instruments are in the microsecond timescale (about 100 μs per TOF spectrum). As IMS of gas phase ions is accomplished in the millisecond range it can be implemented as a third dimension of separation in between chromatographic separation and mass analysis in a loss-free way[8]. In 2015, Meier et al. introduced a novel scan mode termed parallel accumulation – serial fragmentation (PASEF) [5, 6] which has been implemented on a trapped ion mobility quadrupole time-of-flight (TIMS-Q-TOF) mass spectrometer[9] synchronizing mobility separation with quadrupole mass selection. Moreover, PASEF incorporates a dual TIMS technology where in-coming ions are accumulated in the front section, while ions in the second TIMS analyzer are sequentially released depending on their ion mobility markedly increasing duty cycle, sensitivity and spectrum acquisition rate in shotgun proteomics experiments. PASEF, initially implemented as DDA-based approach, was recently combined also with DIA [9], and further refined to optimize the ion mobility and mass ranges for best precursor coverage or sensitivity [10, 11]. Within each TIMS cycle, diaPASEF strategies typically apply fixed, non-overlapping quadrupole isolation windows for ion selection in the m/z domain which are then synchronized with mobility separation (i.e., the release of the target ions from the TIMS device). To achieve sufficient selectivity for the analysis of highly complex proteome samples, each TIMS scan comprises a comparatively low number of non-overlapping quadrupole isolation windows (typically two to five). This approach negatively impacts sensitivity as not all precursors within the mass range of interest are selected for fragmentation during a TIMS scan. Narrowing down each window in the mobility dimension 1/K0 range (0.03-0.045) while concomitantly increasing quadrupole isolation window width as suggested by slicePASEF reduces the number of TIMS cycles required for precursor fragmentation and enhances sensitivity [12]. Recently, Amodei et al. demonstrated that a marked improvement in precursor selectivity can be obtained by offsetting successive cycles of isolation windows with respect to each other [13]. As the only change to the acquisition method is shifting the location of the isolation windows in alternating scan cycles, referred to as “overlapping” window mode, the concomitant increase in precursor precision has no negative impact on the duty cycle of the instrument. Alternative approaches that increase precursor selectivity are SONAR [14] and scanningSWATH [15]. Here, a “sliding” quadrupole scheme is used that scans over the mass range of interest thereby transmitting ions in sequence facilitating fragment-precursor ion assignment and increasing duty cycle as compared to fixed window schemes. In the present work, we introduce and characterize a novel DIA strategy/scan mode that integrates the concepts and benefits of diaPASEF, scanning quadrupole and overlapping quadrupole window acquisition. Applying mobility-specific micro-encoding of overlapping quadrupole window positions optimally covers the ion population in the IM – m/z plane, thereby **M**axmizing **I**nformation content in **DIA**-PASEF acquisitions, which we termed midiaPASEF. We established and integrated bioinformatic tools to enable automated data processing of resulting high complexity datasets. Additionally, we demonstrate that the use of overlapping windows provides a 2.4-fold increase in fragment ion sensitivity as compared to the “classical” diaPASEF approach while improving the precision of precursor-fragment relationships to < 2 Th. Lastly, we show that resulting deconvoluted MIDIA-MSMS fragment ion spectra display DDA-like quality and achieve a fragment ion mass accuracy below 10 ppm.

## 2 Material and Methods

### 2.1 Cell Culture

The human cervix carcinoma cell line HeLa was obtained from the German Resource Centre for Biological Material (DSMZ). Cells were cultured in Iscove’s Modified Dulbecco’s Medium (IMDM; PAN Biotech, Aidenbach, Germany) supplemented with 10% (v/v) fetal calf serum (FCS; Thermo Fisher Scientific (Invitrogen), Waltham, MA), 1% (v/v) L-glutamine (Carl Roth), and 1% (v/v) sodium pyruvate (Serva, Heidelberg, Germany) at 37°C in a 5% CO2 environment and harvested at 70% confluence. Cells were washed once with phosphate buffered saline (PBS; Carl Roth) prior to detaching from the culture flasks. After detaching with 0.05% Trypsin-EDTA solution (Sigma-Aldrich), cells were transferred into centrifugal tubes and washed three times with PBS. Cell pellets were stored at -80°C until further processing.

### 2.2 Protein extraction and preparation of whole proteome samples

HeLa cells were lysed using an urea-based lysis buffer (7 M urea, 2 M thiourea, 5 mM dithiothreitol (DTT), 2% (w/v) CHAPS). Cell lysis was further promoted by sonication at 4°C for 15 min using a Bioruptor device (Diagenode, Liège, Belgium). After cell lysis, protein concentration was determined using the Pierce 660 nm protein assay (Thermo Fisher Scientific) according the manufacturer’
ss protocol and samples were processed using filter-aided sample preparation (FASP) as detailed before[16, 17]. In brief, lysates were loaded onto spin filter columns (Nanosep centrifugal devices with Omega membrane, 30 kDa MWCO; Pall, Port Washington, NY) and washed three times with buffer containing 8 M urea. Afterwards, proteins were reduced and alkylated using DTT and iodoacetamide (IAA), respectively. After alkylation, excess IAA was quenched by the addition of DTT. Afterwards, buffer was exchanged washing the membrane three times with 50 mM NH_4_HCO_3_ and proteins digested overnight. Overnight digestion was conducted at 37°C using trypsin (Trypsin Gold, Promega, Madison, WI) at an enzyme-to-protein ratio of 1:50 (w/w). After proteolytic digestion, peptides were recovered by centrifugation and two additional washes with 50 mM NH_4_HCO_3_. After combining peptide flow-throughs, samples were acidified with trifluoroacetic acid (TFA) to a final concentration of 1% (v/v) TFA and lyophilized. Lyophilized peptides were reconstituted in 0.1% (v/v) formic acid (FA) for LC-MS analysis.

### 2.3 Preparation of phosphopeptide samples

Mice (C57BL/6) were sacrificed by CO_2_ asphyxiation. After decapitation, the brain was dissected, immediately frozen using liquid nitrogen and stored at -80°C. The animal experiments were conducted in accordance with national laws and approved by the local authorities. Tissue was grinded in liquid nitrogen using a mortar and pestle. Afterwards, proteins were extracted from the tissue powder adding a urea-based lysis buffer (8 M urea, 2 M thiourea in 100 mM NH_4_HCO_3_, pH 7.4). To further promote lysis, samples were sonicated for 15 min (30 s on/30 s off) at 4 °C in a Bioruptor device (Diagenode, Belgium). After reduction and alkylation with DTT and IAA, proteins were digested overnight at 32 °C using trypsin (Pierce TPCK-Trypsin, Thermo Scientific) an enzyme-to-protein ratio of 1:25 (w/w). After overnight digestion, peptides were desalted using SepPak tC18 100 mg cartridges (Waters Corporation) and lyophilized. Phosphopeptide enrichment was performed using preloaded TiO2 spin-tips (3 mg TiO_2_, 200 μL tips, GL Sciences, Tokyo, Japan). The tips were conditioned at room temperature (RT) by centrifugation (3,000 x g, 2 min) passing through 20 μL of wash buffer (80 % (v/v) acetonitrile (ACN), 0.4 % (v/v) TFA) followed by 20 μL of loading buffer (57 % (v/v) ACN, 0.3 % (v/v) TFA, 40 % (v/v) lactic acid) applying the same settings for centrifugation. The peptides were resuspended in 150 μL loading buffer, loaded onto the spin-tips and centrifuged (1,000 x g, 10 min, RT). The flow-through was re-applied and centrifuged with same settings. Bound phosphopeptides were first washed with 20 μL loading buffer followed by three wash steps with 20 μL wash buffer (all at 3,000 x g, 2 min, RT). Purified phosphopeptides were then eluted by centrifugation (1,000 x g, 10 min, RT) adding first 50 μL of 1.5 % (v/v) NH4OH followed by 50 μL of 5 % (v/v) pyrrolidine. Eluted phosphopeptides were acidified adding 100 μL 2.5 % (v/v) TFA and desalted using Pierce graphite spin-columns (Thermo Scientific) following the manufacturer’s protocol. After elution and lyophilization, the phosphopeptides were reconstituted in 20 μL 0.1 % (v/v) FA for LC–MS analysis.

### 2.4 Liquid-chromatography mass spectrometry (LC-MS)

For the LC-MS analysis of HeLa dilution series, reconstituted HeLa peptides were separated on a nanoElute LC system (Bruker Corporation, Billerica, MA, USA) at 850 nL/min using a reversed phase C18 column (PepSep, 25 cm x 150 μm 1.5 μm, Bruker Corporation) attached to a CaptiveSpray Emitter (20 μm, Bruker Corporation). The column was heated to 40°C. Mobile phase A was 0.1% FA (v/v) in water and mobile phase B 0.1% FA (v/v) in ACN. Peptides were loaded onto the column in direct injection mode at 800 bar and were separated running a linear gradient from 2% to 38% mobile phase B over 35.5 min. Afterwards, the column was rinsed at 95% B resulting in a total method time of 41 min. Phosphopeptides as well as MHC class I ligands were analyzed using a C18 Aurora UHPLC emitter column (25 cm x 75 μm 1.6 μm, IonOpticks), which was heated to 50°C. Peptides were loaded onto the column in direct injection mode at 600 bar and separated running a linear gradient from 2% to 37% mobile phase B over 39 min at a flow rate of 400 nL/min. Afterwards, column was rinsed for 5 min at 95% B. Eluting peptides were analyzed in positive mode ESI-MS on a timsTOF Pro 2 mass spectrometer (Bruker Corporation) modified to enable acquisition of midia-PASEF mode. Data were acquired using parallel accumulation serial fragmentation (PASEF) enhanced data-dependent (DDA) and data-independent acquisition mode (DIA) [9, 18] as well as our novel acquisition scheme, midia-PASEF with other parameters set to values previously optimized for these acquisition methods and samples. For the comparison of the different acquisition schemes, the dual TIMS was operated at a fixed duty cycle close to 100% using equal accumulation and ramp times of 100 ms each spanning a mobility range from 1/K_0_=0.6Vscm^-2^ to 1.6Vscm^-2^. For the diaPASEF analyses, we defined 44×25 Th isolation windows from m/z 300 to 1,165 resulting in twenty diaPASEF TIMS scans per acquisition cycle and a total cycle time of 2.2 s (to match the duty cycle of the corresponding MIDIA runs). The collision energy was ramped linearly as a function of the inverse mobility from 59eV at 1/K_0_=1.3Vscm^-2^ to 20eV at 1/K_0_=0.85Vscm^-2^. The DDA-PASEF mode comprised ten PASEF scans per topN acquisition cycle [9]. Singly charged precursors were excluded from fragmentation by their position in the m/z–ion mobility plane. For midia-PASEF acquisition, we programmed twenty scans, i.e. MIDIA frames, with quadrupole isolation window widths of 36 Th (after quadrupole calibration for scanning mode) resulting in an overall cycle time of around 2.2 s. Quadrupole isolation window positions were shifted 12 Th (as compared to the previous frame) during acquisition resulting in an overlap of 24 Th between two neighbouring frames. In the MIDIA method, precursor mass ranges covered m/z = 121 - 1565 Th. The collision energy was ramped linearly as a function of the mobility from 70 eV at 1/K_0_=1.56Vscm^-2^ to 20 eV at 1/K_0_=0.60Vscm^-2^. A fixed value of 10 eV was set for experiments, where no peptide fragmentation was induced and unfragmented precursors were recorded.

### 2.5 Data processing

The DDA raw files were processed by PEAKS X Pro v10.6 (BSI, Canada). Phosphopeptide samples were searched using a custom compiled database containing UniprotKB/Swissprot entries of the mouse reference proteome (UniPro-tKB release 2022_04, 17,107 entries) and a list of common contaminants. MHC samples were searched against a human reference proteome database (UniProtKB release 2022_04, 20,361 entries). Precursor and fragment ion tolerance were set to 15 ppm and 0.03 Da, respectively. In case of the phosphoproteome samples, trypsin was set as enzyme for digestion allowing up to two missed cleavages. Carbamidomethylation at cysteines was set as fixed modification. Methionine oxidation, N-term acetylation as well as phosphorylation on serine, threonine and tyrosine were set as variable modifications allowing a maximum of five variable modifications per peptide. The following parameters were chosen to search the MHC dataset: No enzyme was specified and digestion mode was set to “unspecific”. Cysteine carbamidomethylation, methionine oxidation and cysteinylation were set as variable modifications allowing a maximum of two variable modifications per peptide. DIA raw data were processed using DIA-NN (v1.8) [19] applying the default parameters for library-free database search. Data were searched against the same databases as specified above, i.e. for DDA analysis (mouse reference proteome: UniProtKB release 2022_04, 17,107 entries and human reference proteome: UniProtKB release 2022_04, 20,361 entries). For peptide identification and in-silico library generation, trypsin was set as protease allowing one missed cleavage. Carbamidomethylation was set as fixed modification. The maximum number of variable modifications was set to zero. In case of the phosphoproteome dataset, the maximum number of variable modifications was set to 3, allowing only UniMod:21 modifications, i.e. mass delta of 79.9663 Da corresponding to phosphorylation at serine, threonine and tyrosine. The peptide length ranged between 7–30 amino acids. The precursor m/z range was set to 300 – 1,800 Th, and the product ion m/z range to 200 – 1,800 Th. As quantification strategy we applied the “any LC (high accuracy)” mode with RT-dependent median-based cross-run normalization enabled. We used the in-build algorithm of DIA-NN to automatically optimize MS2 and MS1 mass accuracies and scan window size. Peptide precursor FDRs were controlled below 1 %.

### 2.6 Machine learning for refinement of database searches

Peaks were detected for both precursor and fragment ions using the four-d feature finder (4DFF) tool developed by Bruker. Since as of today, 4DFF is not yet adapted to deal with the MIDIA acquisition scheme directly, this was made possible by custom partitioning of the dataset. Subsequently, we generated a set of candidate relations by coarsely matching all precursor peaks with all fragment peaks based on a cutoff distance in retention time and scan dimension of 2.1 sec and 10 scans, respectively. For refinement of relationships, these candidates were then labeled either one or zero, depending on a precursor-fragment pair being part of a PEAKS identified peptide. Features containing information about peak distributions were either provided by 4DFF or calculated from the given statistics. A precursor fragment relation is ultimately described by a feature vector of 15 values containing information about pairwise distances in retention time and scan as well as peak distributions along the recorded dimensions.

From that, a training set was constructed containing a balanced number of positive and negative examples by down-sampling of negative examples. A random train-validation-test split of ≈ 80% training and ≈ 10% validation and test was performed. This resulted in a total of 4,889,621 examples for training and 1,222,406 examples for validation and testing. A gradient boosted tree [20], was trained with default parameters on the training portion of the dataset using xgboost version 1.7.2 for Python. It learned to decide for a given relation between a precursor and a fragment ion whether or not it should be kept for downstream analysis. After training the tree model, the validation set was used to find an optimal cutoff-value in form of the lowest probability output at which a relation should still be considered. Cutoff values were chosen such that the model-recall (number of selected relations divided by all positively labeled relations) reached defined values between 90 and 99%. This classifier and probability cutoff was then used to refine the set of coarsely generated precursor-fragment relations, which were exported to .mgf or .ms2 files for database search using an in-house developed module written in the Python programming language.

## 3 Results

### 3.1 Hardware development

In the present study, we present and characterize a novel scan mode[21], midiaPASEF, that combines the concepts of diaPASEF[18] scanning quadrupole [15, 14] and overlapping quadrupole window acquisition[13] (see Figure 1). To acquire data in midiaPASEF mode, a precise control of the quadrupole positioning in a TOF-scan specific manner is required. To enable this, we added the quadrupole control to the field programmable gate array (FPGA) based part of the instrument controller that allows to assign specific quadrupole positions (defined by center and intended width of the isolation window) for each individual TOF scan in each MIDIA acquisition frame. For a typical 100 ms PASEF accumulation/scan time, and a pusher cycle time of 110 μs, this translates into 927 programmable quadrupole isolation window positions for each 100 ms MIDIA frame. Due to the time constants of the RC elements of the analogue part of electronics and ion optics, this high frequency of digital quadrupole control steps results in a continuously moving quadrupole isolation window (scanning quadrupole). To optimally cover the precursor ion cloud in the m/z-1/K_0_ space, we selected an isolation parallelogram with a fixed width of 264 Th at each 1/K_0_ value, covering a total mass range from 121 to 1535 Th. The position and steepness of the isolation parallelogram was optimized to provide optimal coverage of the analyte space of tryptic peptides from HeLa. In midiaPASEF acquisition, this parallelogram is covered by 20 diagonals (i.e. MIDIA frames) of 36 Th quadrupole isolation width each and 24 Th overlap between adjacent diagonals, resulting in a shift of 12 Th between two subsequent MIDIA frames. The isolation pattern of a single MIDIA frame is depicted in Figure 2 A, lower panel, the overall parallelogram is shown in Supplementary Figure 1A. The combination of scanning quadrupole with an overlapping window scheme (Supplementary Figure 1B) results in multiple passes of quadrupole isolation edges for each precursor ion (Figure 1 B,C) and leads to a characteristic transmission matrix specific for each precursor m/z-value, TOF-scan (1/K_0_) and MIDIA frame. Within one full MIDIA cycle, each precursor is covered by three subsequent MIDIA frames at each distinct 1/K_0_-range (i.e. scan region), resulting in a 3-fold increase in sensitivity compared to non-overlapping windows with the same offset between frames. This acquisition scheme defines unique transmission patterns, which we termed “MIDIA fingerprints” that are highly specific for the respective precursor m/z and 1/K_0_ combination. (Figure 1D). To generate high-resolution fragment spectra, ion populations transmitted by the quadrupole are fragmented in the collision cell and subsequently analyzed with a TOF resolution of 50,000, thereby providing the basis for high mass accuracy fragment ion spectra.

**Figure 1:**
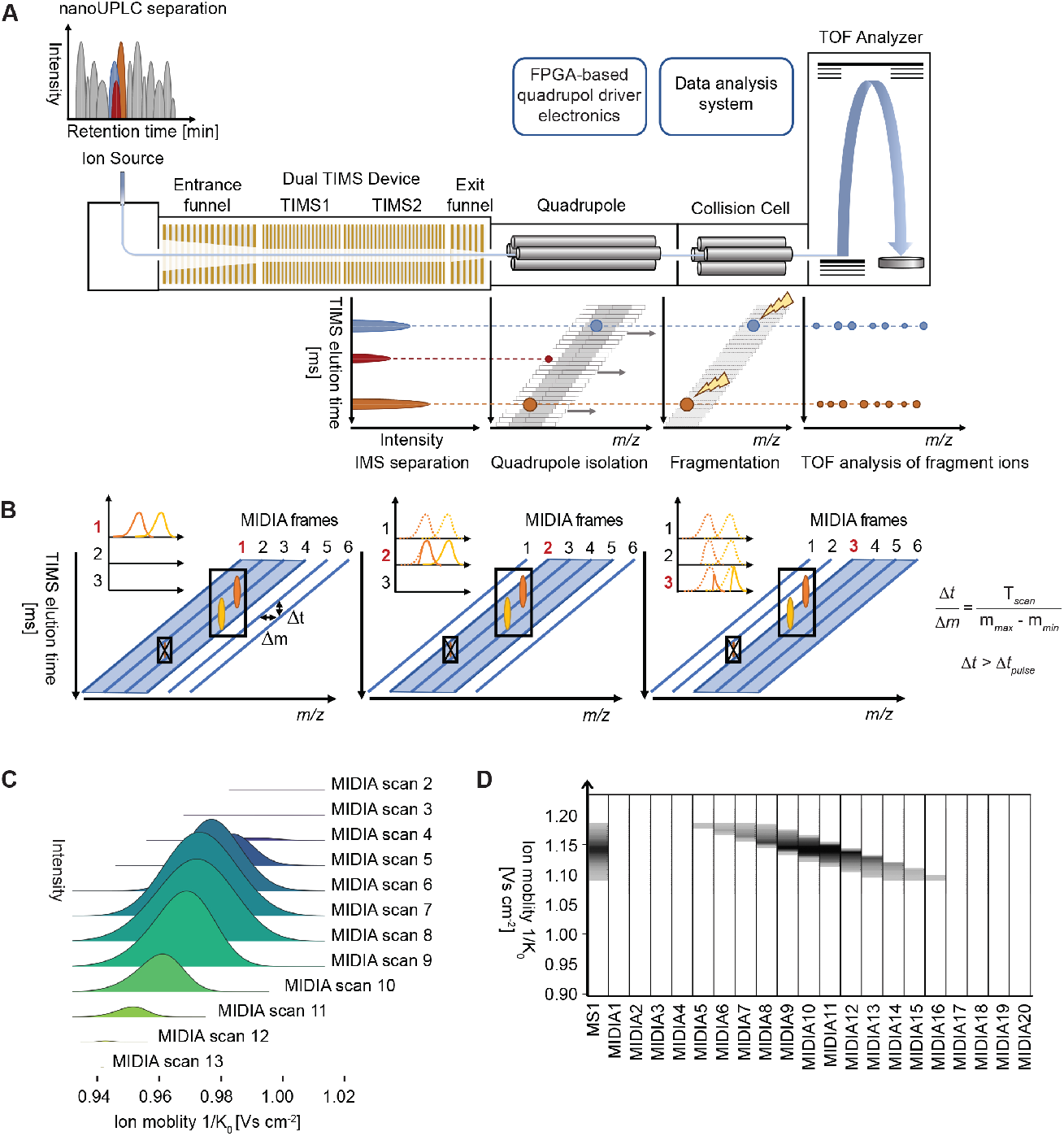
Overview of the midiaPASEF acquisition concept. A) Schematic view of the timsTOF Pro 2 instrument and midiaPASEF acquisition mode. Peptides eluting from the nanoUPLC are ionized and enter the dual TIMS device. In the first TIMS analyzer ions are accumulated, while a second ion package is released in parallel from the second TIMS device according to ion mobility. For precursor fragmentation, the quadrupole is operated in scanning mode and synchronized to the TIMS elution of ions of interest covering the full 1/K_0_ range within one MIDIA frame. For a TIMS cycle time of 100 ms and a pusher cycle time of 110 μs this translates into 927 programmable quadrupole isolation windows. To enable such a fast, continuous quadrupole movement, the digital analog converters of the quadrupole RF-amplitude, the bias-DC, and the resolution-DC are routed through a built-in FPGA. Ion populations transmitted by the quadrupole are fragmented in the collision cell and subsequently analyzed in the TOF region. B) Principle of the overlapping windows scanning scheme in midiaPASEF. To optimally cover the precursor ion cloud and to facilitate precursor-fragment-ion mapping, quadrupole isolation widths in each frame are set to 36 Th and shifted by 12 Th in the subsequent MIDIA frame resulting in an overlap of 24 Th between two neighbouring MIDIA frames. The combination of scanning quadrupole with an overlapping window scheme leads to a transmission matrix specific for each m/z-value, scan (1/K_0_) and MIDIA frame. C) MIDIA transmission profile across multiple MIDIA frames defined by overlapping window acquisition result in multiple passes of quadrupole edges for each precursor ion. D) Within one full MIDIA cycle, precursors are covered by three subsequent MIDIA frames at a distinct 1/K_0_-range (i.e. scan region), resulting in a 3-fold increase in sensitivity compared to non-overlapping windows and defining unique “MIDIA fingerprints” that are highly specific for the respective precursor m/z - 1/K_0_ combination.

**Figure 2:**
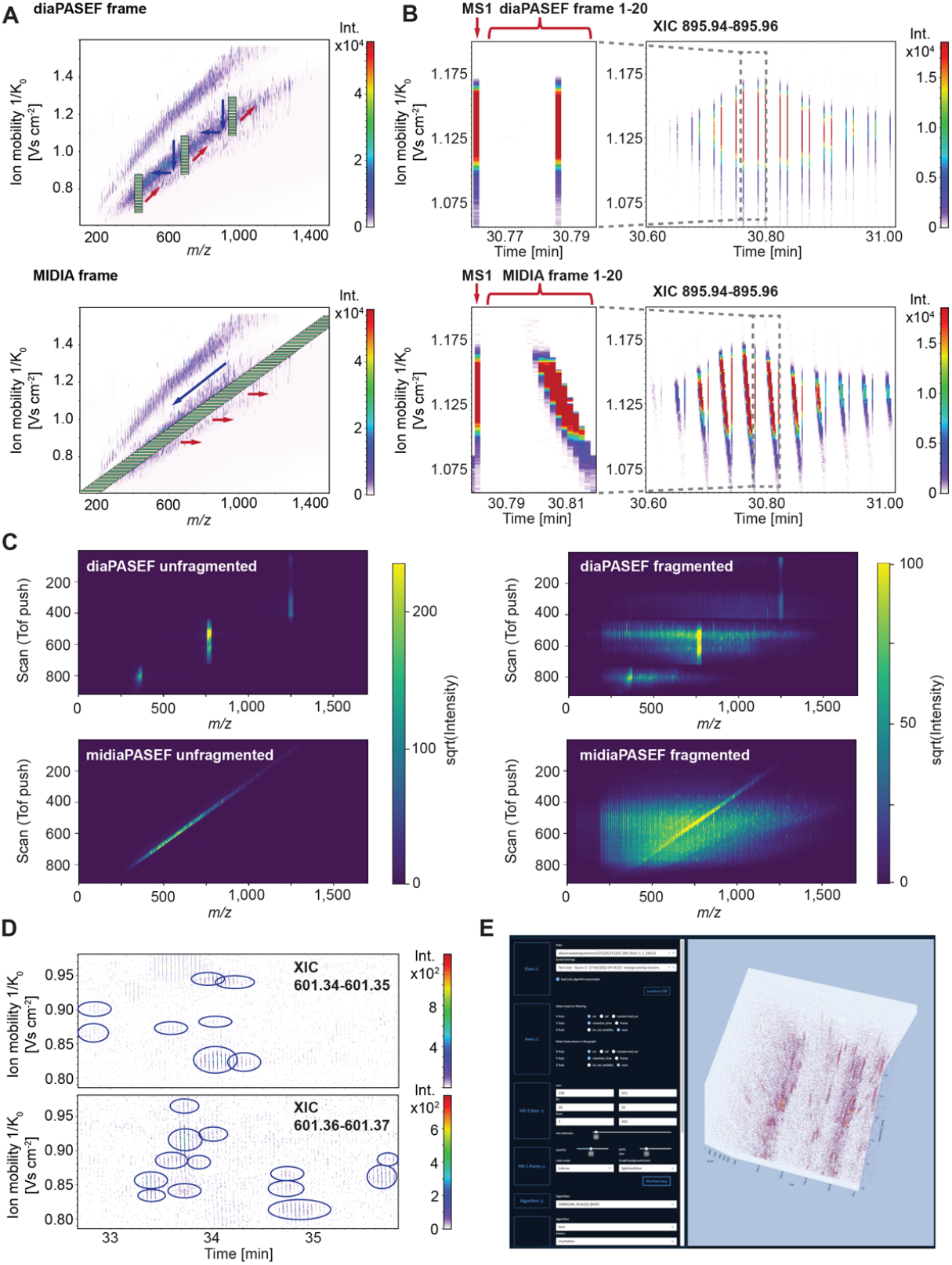
Comparison of diaPASEF and midiaPASEF scan modes. To compare the different acquisition modes, 100 ng of tryptic HeLa lysate was separated and analyzed by LC-MS using a 40 min gradient. A) Acquisition scheme of diaPASEF (upper panel) and midiaPASEF (lower panel). Depicted is a single frame. Red arrows indicate the movement of quadrupole isolation windows in subsequent frame(s) and blue arrows the quadrupole movement within a single diaPASEF or midiaPASEF frame. B) Comparison of precursor transmission profiles for diaPASEF and midiaPASEF (precursor m/z 895.95) at low collision energies (no fragmentation). During diaPASEF acquisition precursors are only selected in a single diaPASEF frame for subsequent fragmentation, whereas precursor ions are isolated in multiple frames in midiaPASEF resulting in a three-fold increase in sensitivity. C) Isolation and fragmentation patterns in diaPASEF and midiaPASEF. In midiaPASEF precursors are fragmented across the full ion mobility range, whereas in diaPASEF mode ions at low mobility regions and at the transition points between PASEF windows are not fragmented. D) midiaPASEF acquisition results in highly complex datasets. XIC of two neighbouring mass ranges (m/z 601.34-601.35 and 601.34-601.35) in the time range between 32.8 min and 35.8 min demonstrate the high complexity of midiaPASEF datasets and observation of the characteristic “MIDIA fingerprints”. E) 3D view of midiaPASEF data in the MIDIAviewer package.

### 3.2 Characterizing quadrupole transmission profiles

For MIDIA type acquisition, the isolation scheme described above translates into an offset in the quadrupole position of 1.2 Th for 100 ms and 2.4 Th for 50 ms between subsequent TOF scans. This in turn results in very high quadrupole “scanning speeds” of approx. 11,000 amu/sec (100 ms) or 22,000 amu/sec (50 ms). As these scanning speeds are outside of the range of the initial quadrupole design specifications, we hypothesized that they might modify the effective shape of the quadrupole transmission window, as the quadrupole set position effectively changes during the transition time of an ion through the quadrupole. Therefore, we evaluated the shape of the effective quadrupole isolation windows using LC-IMS-Q-TOF runs of a tryptic digest of HeLa with a low collision energy of 10 eV, which does not induce fragmentation of peptides. The use of unfragmented data allows easy mapping of signals between MS1 and MIDIA space as the precursor m/z remains unchanged. By integrating datapoints over an entire LCMS run, precise transmission coefficients for each m/z can be calculated as a tensor defined by MIDIA frame, TOF scan and m/z. Hereby, we obtained transmission profiles for each MIDIA frame and scan position. Transmission profiles were fitted and the initial fits were integrated over the entire acquisition range, leading to a transmission matrix specific for each MIDIA frame and TOF-scan. Our analyses indicated that both shape and position of the isolation window are significantly different from programmed values or the respective values obtained from standard diaPASEF acquisitions. Observed quadrupole transmission window shapes are narrower (as defined by the distance of the 50 % transmission points) and display a marked broadening of the isolation flanks. This effect can be explained by the very fast ramping speeds, as the quadrupole changes position in a time frame that is shorter than the transit time of the ions through the quadrupole. To compensate for this effect, we developed a quadrupole calibration procedure, which typically limits the difference between programmed and fitted quadrupole windows to < 2 Th (Data not shown).

### 3.3 Comparison between diaPASEF and MIDIA acquisition modes

Having precisely determined the quadrupole transmission properties, we compared the transmission pattern of ions between diaPASEF and MIDIA modes. In diaPASEF, the quadrupole isolation window (typically 25 Th) is kept constant for a certain 1/K_0_ range and is then switched to the next position (Figure 2A, top panel). This results in each precursor ion to be transmitted typically in only a single of the 16-20 diaPASEF frames, thus limiting the transmission efficiency (Figure 2B, top panel). In contrast, midiaPASEF acquisition changes the quadrupole transmission window with each TOF scan, thereby adapting to the shape of the precursor ion cloud (Figure 2A, lower panel). midiaPASEF acquisition results in a typical diagonal transmission pattern for unfragmented data (Figure 2C). Applying collision energy results in the efficient generation of fragment ions in a TOF-scan and quadrupole window specific manner (Figure 2C, bottom right panel).

The 36 Th wide transmission windows are shifted by 12 Th between adjacent MIDIA frames, leading to transmission of each precursor ion in three consecutive MIDIA frames, thereby increasing transmission efficiency 3-fold over a non-overlapping acquisition scheme with the same offset (Figure 2B, lower panel). Furthermore, the specific design of the quadrupole isolation scheme leads to characteristic and highly specific transmission patters, i.e. MIDIA fingerprints, that can be observed within each MIDIA acquisition cycle (1 MS1 frame, followed by 20 MIDIA frames). An example for such a MIDIA fingerprint is displayed for a doubly charged precursor ion with an m/z of 895.95 in Figure 2B (lower panel).

The resulting raw data are highly complex and very rich in information content, as exemplified by fragment ion clusters detected in a tryptic digest of HeLa in two adjacent 10 mDa mass ranges (Figure 2D). To enable a visualization and graphical explorative analysis of the MIDIA datasets, we developed the MIDIAviewer Tool (Figure 2E, which is based on OpenTIMS [22] and allows to visualize the multidimensional datasets.

### 3.4 Precursor ion prediction precision

The characteristic information-rich MIDIA fingerprints (i.e. transmission patterns of ions) are observable even for very low intensity ions as exemplified by the observed MIDIA fingerprints for the 1^st^, and 7^th^ isotope of a peptide precursor ion (m/z = 488.28, z = 2^+^). Notably, the theoretical isotopic abundance of the 7^th^ isotope of the peptide (m/z = 491.28) is about 10,000-fold lower than the first isotopic peak, illustrating the applicability of midia-PASEF even for very low-abundant ions. (Figure 3A). The information content encoded in the MIDIA fingerprints of each fragment ion provides the basis for the highly specific prediction of the respective m/z position of the corresponding precursor ion. Towards this purpose, we developed and tested several machine-learning algorithms that output score distributions for possible precursor positions for each observed fragment ion based on its MIDIA fingerprint (Figure 3B). To evaluate the prediction performance on the level of an entire dataset, we again used unfragmented datasets acquired at low collision energy from tryptic digests of HeLa proteome. In this case, as no fragmentation occurs, the m/z of each MS2 ion corresponds to its “precursor ion m/z”. Analyzing more than 200,000 ion clusters across over four orders of magnitude dynamic range, we confirmed a prediction specificity of < 2 Th for > 90 % of observed MS2 ions for the best performing precursor prediction algorithms (Figure 3C). Next, we evaluated the performance of our precursor prediction algorithms for midiaPASEF data for fragment ions. As an example we picked several fragment ions across the full intensity range, i.e. from high to low abundant ions, of the peptide SLETEMSALQLQVTER that had been identified by PEAKS from a deconvoluted midiaPASEF spectrum (Figure 3D) across the intensity range. For high intensity fragment ions such as the y_7_ ion (m/z = 873.48) the respective MIDIA fingerprints consist of almost complete observation matrices (Figure 3E). This holds true also for fragment ions in the medium range, such as the y_9_ (m/z = 1057.6) and also the b_6_ ion (m/z = 674.30). Less complete matrices are observed for lower intensity fragments, such as the y_2_ ion (m/z = 304.14, 6.5% relative intensity) and rather sparse MIDIA fingerprints are obtained for very low intensity fragments, e.g. ammonia losses, such as y_6_–NH_3_ (m/z = 728.40). Ions in the lowest intensity range e.g. y_10_-NH_3_ (m/z = 1127.62, 0.9% relative intensity) are at the edge of the detection limit. Notably, precursor prediction remains accurate even for very low intensity fragment ions with as little as 8-10 ion signal observations (Figure 3F).

**Figure 3:**
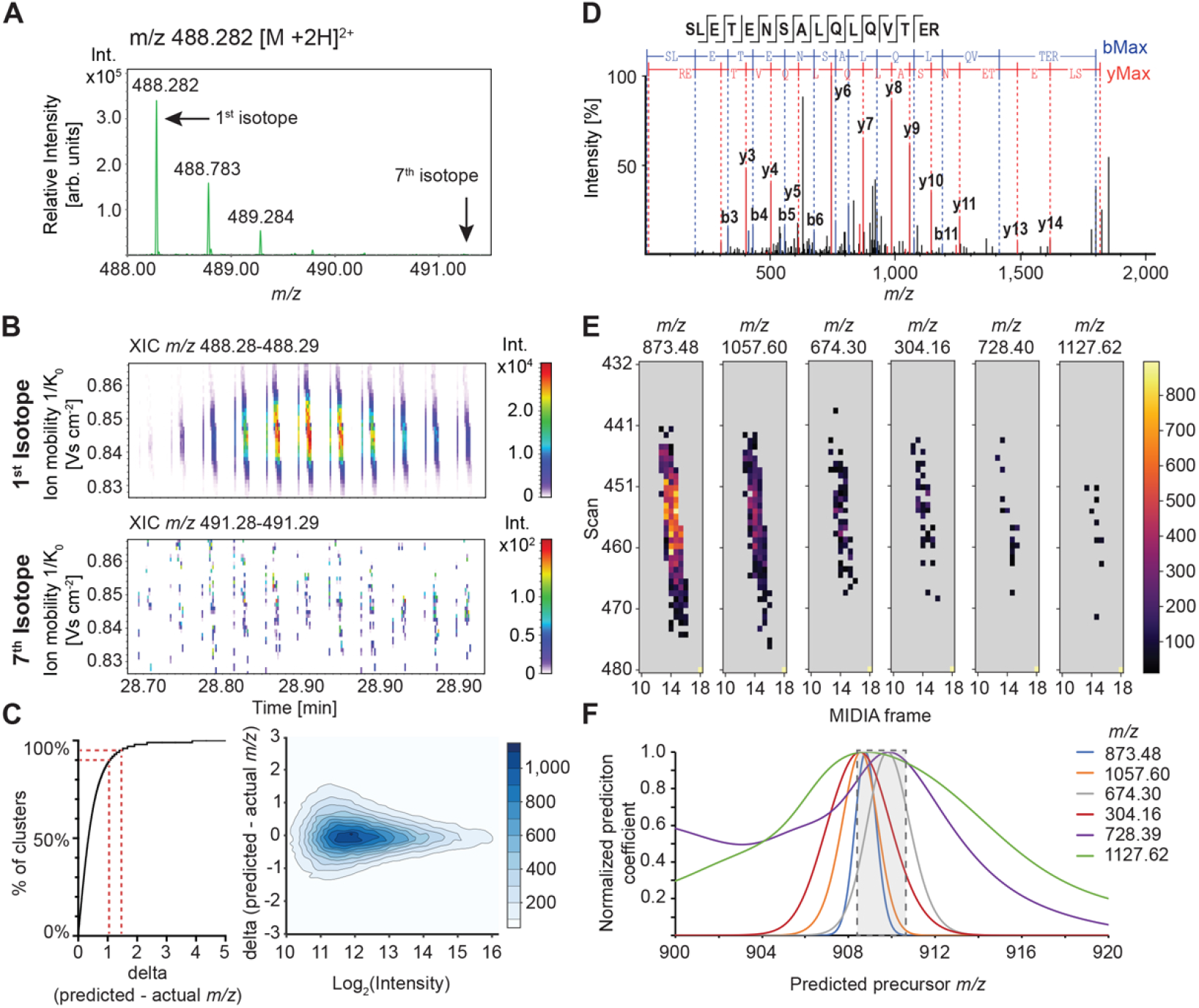
MIDIA fingerprints are information rich and enable precise prediction of precursor m/z values across the full intensity range. A) MS1 spectrum of a peptide MS1 ion (m/z 488.282, 2+) B) Comparison of midia-PASEF transmission patterns (“MIDIA fingerprints”) of the first (m/z 488.28) and seventh (m/z 491.25) isotopic peak. MIDIA fingerprints are observable across several orders of magnitude in a single acquisition cycle and integrate specific precursor position information associated with each TOF scan and thus enable precise prediction of precursor m/z positions for low intensity ions. C) Evaluation of precursor prediction performance on unfragmented (low collision energy) data. 300 ng of HeLa digest were analyzed by midiaPASEF without inducing fragmentation. Precursor positions were predicted using machine-learning based methods and the refined prediction error plotted as a function of the cluster intensity. On this dataset comprising >200,000 ion observations, 80% of precursor positions can be predicted with a deviation of <1 Th, and 95% with a precision of <2 Th, achieving a precursor specificity comparable to DDA across the full dynamic range. D) Example deconvoluted MIDIA-MSMS spectrum of peptide SLETEMSALQLQVTER identified by PEAKS X Pro database search. E) MIDIA fingerprints of selected fragment ions of peptide SLETEMSALQLQVTER integrate specific precursor position information from all TOF spectra acquired across the full elution profile. F) Corresponding calculated precursor prediction probability profiles for selected fragment ions of peptide SLETEMSALQLQVTER enable precise prediction of precursor m/z positions even for very low intensity fragment ions.

### 3.5 Data processing pipeline

To enable automated and reproducible processing of midiaPASEF dataset, we developed the MIDIAID pipeline which is schematically depicted in Figure 4A. MIDIAID is based on Snakemake[23] and incorporates all steps to generate searchable deconvoluted MIDIA-MSMS .mgf or .ms2 files from midiaPASEF raw data. Initially, precursor and fragment level rawdata are pre-processed and clustered using a custom adapted version of the Bruker 4D feature finder (4DFF) and identified non-deisotoped ion clusters are exported to HDF file containers together with their calculated statistics.

**Figure 4:**
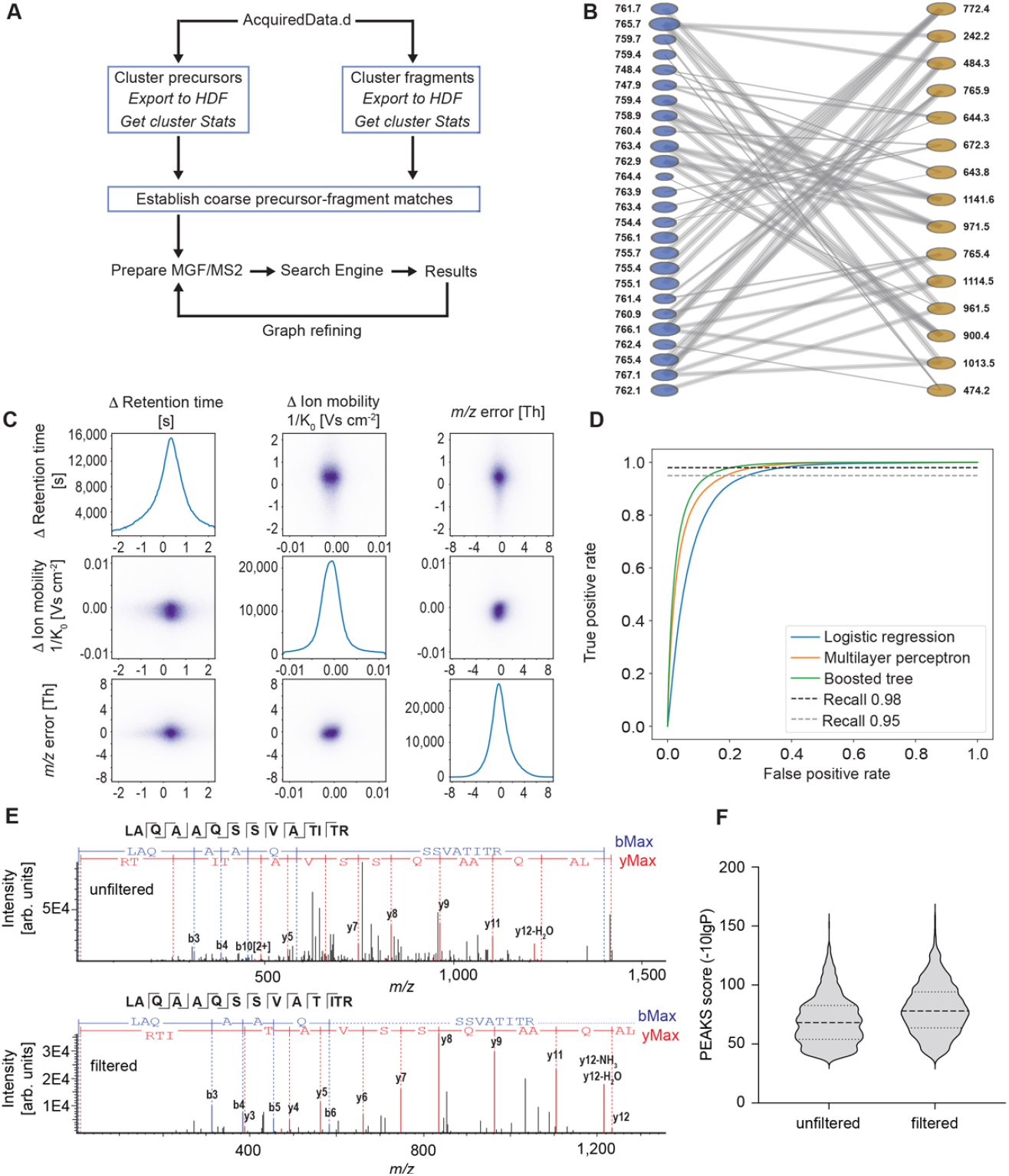
Data evaluation workflow to generate deconvoluted MIDIA-MSMS spectra. A) schematic of the MIDIAID data analysis pipeline. Acquired raw data are clustered using a custom modified version of Bruker 4DFF. Identified ion clusters on precursor and fragment level are exported to HDF and respective cluster statistics are calculated. Subsequently, coarse precursor-fragment relationships are defined based on 3-dimensional distances (frame, scan, predicted precursor m/z). Resulting precursor-fragment relations are exported as MIDIA-MSMS spectra and submitted to first round database search. In an iterative process, precursor-fragment relationships are refined by machine learning and re-submitted to database search. B) midiaPASEF data are stored as bipartite graphs representing MIDIA precursor-fragment relationships. C) 2-dimensional scatterplots of refined key parameters defining precursor-fragment relationships and distances. D) Performance of multiple machine learning models to refine precursor-fragment relationships based on first-round database search results. E) Effects of machine-learning based refining of precursor-fragment relationships for an example peptide (GGVVLKEDALPGQK) from a HeLa tryptic digest (30 min gradient). Data were searched in PEAKS X Pro and fragment ions annotated in red (y-ion series) and blue (b-ion series). Non-annotated ions in the spectra are shown in grey. F) Score distributions of PEAKS X Pro search results of identified HeLa tryptic peptides of initial (unfiltered) and refined (filtered) deconvoluted MIDIA-MSMS spectra.

### 3.6 Bipartite graphs as universal representations of MIDIA precursor-fragment relations

Subsequently, coarse precursor-fragment ion relationships are established based on multidimensional distances in retention time, ion mobility and differences between predicted precursor ion masses of fragment ions and the m/z values of precursor ions. These relationships are stored in the format of bipartite graphs (Figure 4B), where MS1 and MS2 nodes contain properties of the respective MS1 and MS2 ions, and edges contain multidimensional distance information. This graph structure enables a comprehensive description of precursor-fragment relationships [24] and is fully suitable to represent precursor-fragment relationships defined by either DDA or DIA acquisition approaches. Using predefined cut-offs based on the graph structure, MIDIAID then prepares first-round .mgf or .ms2 files for initial database search using established DDA-database search engines including Mascot, PEAKS, Fragpipe[25] and PaSER/ProLucid. Subsequently, first-round database results are filtered at 1 % FDR on peptide level, imported and mapped to the MIDIA-graph. This mapping process results in a set of validated edges in the graph, which are a subset of the total graph. Figure 4D depicts a sub-selection of refined distance characteristics of validated edges in the graph confirmed by PEAKS database search and illustrates that coordinates of MS1 and MS2 ions can be matched with a precision of RT <0.5 sec, 1/K_0_ < 0.005 1/K_0_ and m/z <2 Th for >80% of the precursor-fragment relationships.

### 3.7 Generation of high quality deconvoluted MIDIA-MSMS spectra

To refine the MIDIA graph for the generation of second-round devonvoluted spectra, we applied advanced machine learning tools, including logistic regression, multilayer perception and boosted tree architectures to refine edge properties. Respective ROC curves for selected architectures are depicted in Figure 4E. Notably, this approach reduces the number of edges (i.e. precursor-product relationships) in the graph by 70-80% at a recall rate of 98%. The effects of the graph refining procedure on MIDIA-MSMS spectrum quality before (left panel) and after refining (right panel) are illustrated in Figure 4F. Thereby, our deconvolution algorithms combine the precise information of precursor ion position prediction with correlation metrics in retention time and ion mobility dimensions using machine learning approaches. This process increases spectral purity and generates almost interference-free deconvoluted MIDIA-MSMS spectra with a significantly reduced proportion of unassigned ions, which might derive from co-eluting precursors. Analyzing score distributions of PEAKS search results obtained from database searches of first and second round .mgf spectra confirmed an increase in the median PEAKS score from 80 to 90 (Figure 4G). This recursive strategy renders the precursor specificity comparable to DDA type acquisitions, as indicated by an exceptional spectral purity of the deconvoluted MIDIA-MSMS spectra resulting in score distributions similar to those of DDA-PASEF data. Confirmingly, MIDIA-MSMS spectra are almost indistinguishable from DDA-based spectra for the human eye (Figure 5A,B).

**Figure 5:**
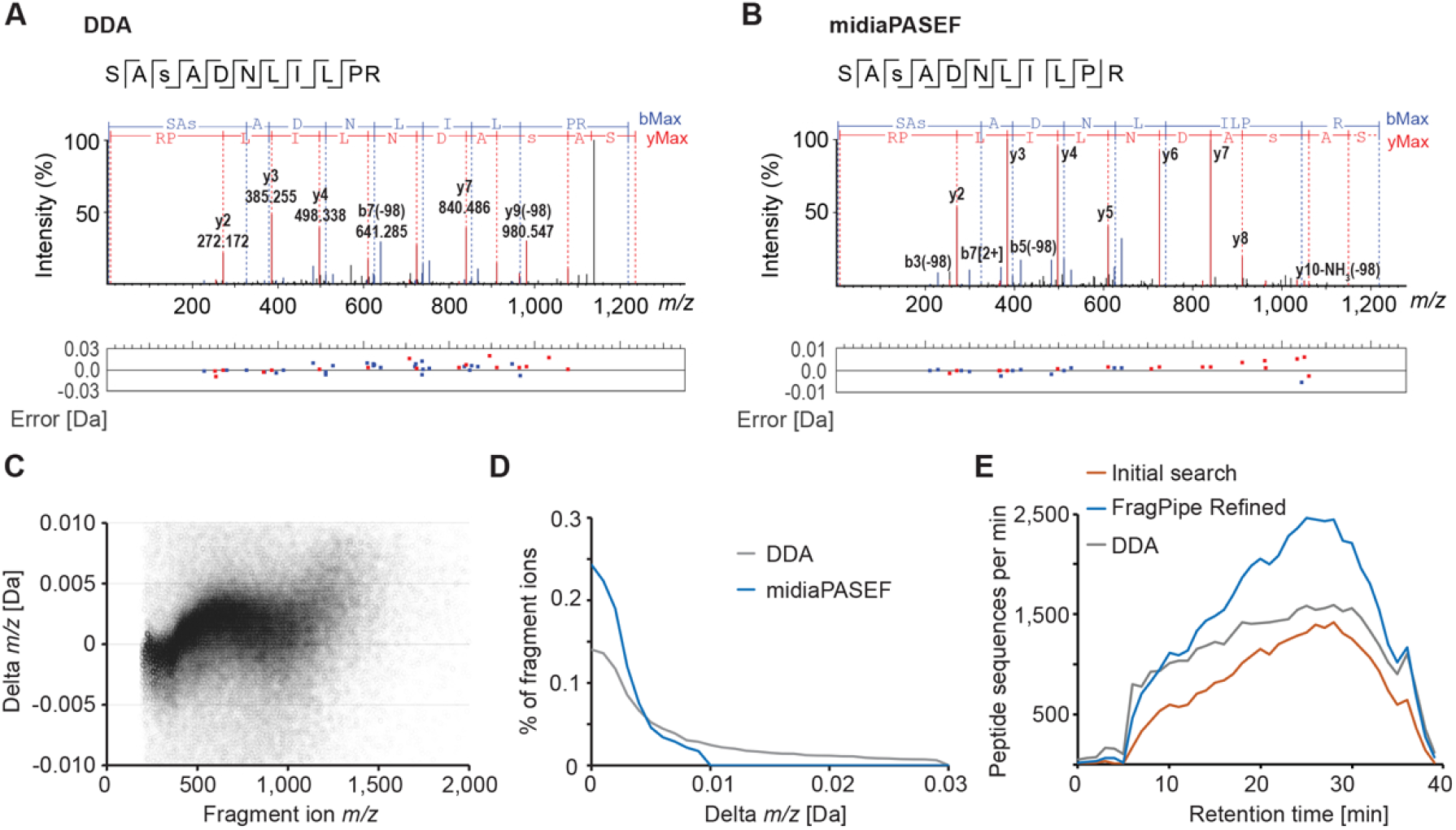
midiaPASEF enables the generation of high quality deconvoluted MSMS spectra. A) DDA and B) MIDIA MSMS fragment ion spectra of the phosphorylated peptide SAsADNLILPR detected in a phosphopeptide mixture derived from mouse brain trytic digest. midiaPASEF acquisition facilitates fragment-precursor assignment resulting in DDA-like spectral quality. This enables the analysis of post-translational modifications, such as phosphorylation, by a DIA-based approach concomitantly providing a MS2 spectral quality that is highly similar to DDA. C) midiaPASEF provides high precision fragment ion m/z values. Deconvoluted MIDIA-MSMS spectra were searched in PEAKS X Pro and resulting mass differences of all identified fragment ions are plotted. D) Analysis of distributions of fragment ion mass deviations of MIDIA and DDA MSMS spectra reveals higher mass accuracy for data collected in midiaPASEF. E) Analysis of database search results of DDA and MIDIA datasets. 300 ng of HeLa tryptic digest was separated on a 150 μm x 150 mm column at 800 nL /min and data acquired in both DDA and MIDIA acquisition modes. DDA data was analyzed in PEAKS. MIDIA data was processed through the MIDIAID pipeline in two iterations. Initial search results reached peptide identification rates of up to 1,400 peptides/min, while database search of the machine-learning refined graph in Fragpipe resulted in the identification of up to 2,487 unique peptide sequences/min (>40 unique peptide sequences per second).

### 3.8 Identification of peptides from MIDIA-MSMS spectra

Having established a machine-learning-based MIDIA graph deconvolution algorithm for the generation of high-quality MIDIA-MSMS fragment spectra, we applied MIDIA towards the analysis of complex proteomic and phosphopeptidomic samples. Deconvoluted MIDIA-MSMS spectra were exported as .mgf files and initially searched in PEAKS. Notably, the PEAKS data processing pipeline includes a *de novo* sequencing step, which consistently resulted in well-annotated spectra (Supplementary Figure 2). This proves – for the first time – that *de novo* sequencing of DIA-derived spectra is feasible on dataset level (Supplementary Figure 3), notably, in the case of midiaPASEF, using a quadrupole selection window width of 36 Th. In addition, MIDIA is fully suitable for the analysis of samples previously highly challenging for DIA, as demonstrated by the analysis of TiO_2_ -enriched phosphopeptides from mouse brain. Example spectra illustrate the quality of the deconvoluted MIDIA-MSMS spectra (Figure 5B), but consistently display a high fragment ion mass accuracy of < 10 mDa. In-depth analysis of fragment mass deviations confirmed this observation on dataset level (Figure 5C). While DDA-PASEF search results displayed a fragment ion mass accuracy of 30 mDa, over 80 % of fragment ions in MIDIA-MSMS spectra displayed a mass deviation below 5 mDa (Figure 5D). The high fragment ion mass accuracy in the 5-10 ppm range results from the design of the overlapping MIDIA diagonals. This increases duty cycle approx. three-fold compared to standard diaPASEF acquisitions and thus ensures robust ion statistics which in turn result in high mass accuracy of detected fragment ions. This high mass accuracy enables database searches with tight fragment ion mass tolerances, which renders midiaPASEF suitable for the comprehensive analysis of most challenging sample types, including phosphoproteomics (Figure 5B) and immunopeptidomics (Supplementary Figure 4). To illustrate the performance of our workflow, we analyzed a tryptic digest of HeLa using a 30 min gradient (41 min runtime) using a 150 μm ID column in both DDA-PASEF and midiaPASEF acquisition modes (Figure 5E). In this analysis, DDA-PASEF acquisition enabled the identification of approx. 30,000 unique peptide sequences, and reached a maximum identification rate of approx. 1,500 unique peptide sequences per minute. First round search results of deconvoluted midiaPASEF datasets resulted in slightly lower numbers, and machine-learning-based refinement of the precursor-fragment relationships in the MIDIA graph in combination with database search in FragPipe enabled the identification of > 2,400 unique peptide sequences per minute in the most complex region of the gradient, which translates into > 40 identified unique peptide sequences per second.

## 4 Discussion

In this manuscript, we introduce the midiaPASEF acquisition method which efficiently maximizes information content in data independent acquisition using overlapping, ion mobility dependent quadrupole windows. Notably, in contrast to established DIA approaches, the MIDIA cycle time does not increase with increased overall mass range. MIDIA diagonal scans cover the entire precursor m/z range and the specific design of the overlapping quadrupole windows enhances transmission of the fragments, as each fragment ion is targeted in 3/20 midiaPASEF frames. This is a 2.4-fold improvement over current standard diaPASEF approaches covering a similar target population of ions, which target each precursor region only in e.g. 1/16 frames. Several implementations of midiaPASEF methods can easily be envisioned to maximize sensitivity, minimize cycle time or maximize duty cycle. The midiaPASEF method presented in this work was optimized for maximum coverage of the precursor ion cloud of multiply charged tryptic peptides and thus uses a total selection parallelogram with a width of 264 Da at each 1/K_0_ value, covered by 20 overlapping midiaPASEF frames. Naturally, several parameters can be optimized to specifically tailor the method for individual applications. For example, a reduction to 10 midiaPASEF frames will reduce the width of the selection parallelogram to 144 Th. This limits precursor ion coverage, but results in a reduction of cycle time to 1.1 sec and a concomitant increase in duty cycle for the ions covered in the selection parallelogram. Further reductions in cycle time can be readily achieved by reducing the TIMS accumulation/separation times. Alternatively, to achieve maximum coverage of the ion space, the width, overlap and position selection windows can be specifically tailored to e.g. optimally analyze phosphopeptidomic or immunopeptidomic samples.

midiaPASEF applies TOF-scan specific encoding of quadrupole windows for transmitting precursors, thereby “imprinting” the quadrupole position as an additional dimension in the data space, similar to the concept of a Q1-dimension as initially discussed by Demichev et al. [19]. Each detected ion is therefore associated not only with frame (RT), scan (ion mobility) and TOF (m/z) information, but also associated with a defined quadrupole selection range. These specific multidimensional transmission properties of MIDIA generate unique characteristic transmission patterns - “MIDIA fingerprints” - for each fragment ion that provide sufficient information to predict the m/z of the respective precursor ion of each detected MS2 feature with a prediction accuracy of typically < 2 Th, similar to gas-phase fractionation small window DIA approaches[26].

Recently, Skowronek et al. described a concept termed “synchro-PASEF” [10]. While the synchroPASEF concept also uses diagonal scanning of the quadrupole as a function of 1/K_0_ to optimally cover the precursor ion cloud, and thus is conceptually similar to midiaPASEF, it lacks the benefits of the overlapping window design. synchroPASEF uses a 24 Da wide quadrupole window and thus requires 11 MS2 frames to cover a 264 Th wide selection parallelogram, resulting in each precursor selected for fragmentation in 1/12 acquisition frames using a 24 Th offset between frames. In contrast, midiaPASEF acquires 20 MS2 frames and selects each precursor in 3/21 acquisition frames at a 12 Th offset, which is essential to define the characteristic MIDIA fingerprints. Notably, midiaPASEF does not require the detection of a “slice event” and enables to precisely predict the precursor positions even for very low abundance fragment ions and thus generates high-quality MIDIA-MSMS spectra at dataset level.

To comprehensively analyze our high complexity midiaPASEF datasets, we developed the MIDIA Graph. This concept is based on the ion networks[24] and fully describes precursor-fragment relationships as edges in a bipartite graph, where detected MS1 and MS2 features are included as nodes and their connections are defined as edges (Figure 3B). Edges are associated with defining properties in multiple dimensions, i.e. differences in retention time, ion mobility and differences between predicted and actual precursor m/z. By trimming the graph at defined cutoff values, deconvoluted MIDIA-MSMS spectra with varying degrees of specificity can be easily generated. In combination with machine-learning based refining of edges in the precursor-fragment ion graph, resulting MIDIA-MSMS spectra achieve the same or even higher specificity compared to a classical quadrupole selection in DDA. Thus, MIDIA-MSMS spectra display fewer of the unassigned peaks observed in real DDA spectra that are likely the result of co-isolation of co-eluting precursors.

We show that midiaPASEF data enables the precise determination of precursor ion positions for all detected fragment ions across the entire dynamic range and show that the MIDIAID pipeline provides DDA-like deconvoluted spectra from midiaPASEF data at a spectral quality suitable even for *de novo* sequencing on dataset level using PEAKS (Supplementary Figure 3).

In conclusion, midiaPASEF enables to reproducibly generate DDA-quality MIDIA-MSMS spectra for precursors across the entire mass range in the selection parallelogram. In contrast to DDA, midiaPASEF is non-stochastic and thus generates detailed detection profiles of each fragment ion in all dimensions, which facilitates the highly specific deconvolution and scoring of precursor-fragment relationships. We envision that these additional layers of information can be used in next generation database search engines to further improve the performance of midiaPASEF. The Snakemake-based MIDIAID analysis pipeline accommodates various approaches for clustering, deisotoping, graph- and .mgf generation. We envision further increase in identification performance by using advanced multidimensional feature detection and clustering approaches. Using machine learning approaches to refine the MIDIA graph based on initial database results, we significantly improved the specificity of precursor-fragment relationships, thereby surpassing the spectral purity of DDA in our deconvoluted MIDIA-MSMS spectra. This also demonstrates that the specific data acquisition in the MIDIA frames as described here does significantly increase the information encoded therein over that encoded in the classical non-overlapping diaPASEF frames. To illustrate the benefit of midiaPASEF for challenging sample types, we have demonstrated the principal applicability to phosphopeptidomic (Figure 5A) and immunopeptidomic (Supplementary Figure 4) samples.

midiaPASEF thus provides all benefits of DIA acquisitions, including efficient ion sampling, high duty cycle and excellent reproducibility. In addition, midiaPASEF allows to generate high-quality deconvoluted MIDIA-MSMS spectra that can be readily exported to various data formats and efficiently analyzed with existing and well-established database search engines that were developed for DDA. Moreover, MIDIA acquisition preserves elution profiles in retention time and ion mobility space, and thus enable to precisely determine the coordinates of the respective ions. In contrast to DDA, fragment ion information is collected across the entire peak, improving ion statistics and thus mass accuracy of fragment ions. Therefore, midiaPASEF-based workflows display excellent potential similar to DDA or small-window DIA approaches[27, 26] for spectral library generation. We envision that the application of midia-PASEF will therefore not be limited to proteomic or peptidomic samples, but can be readily optimized for the analysis of peptides, lipids, metabolites or other small molecules in the mass range between 50 and 5,000 Da.

## 5 Acknowledgements

We thank Christina Jung for excellent sample preparation. This work was funded by the German Ministry of Education and Research (BMBF), as part of the National Research Node “Mass spectrometry in Systems Medicine” (MSCoreSys), under grant agreements 031L0217A (to S.T.) and 031L0217B (to A.H.). and the German Research Foundation (DFG, Deutsche Forschungsgemeinschaft), grant numbers SFB 1292-Q1 and TE599/8-1 to S.T., SFB 1292-TP11 and DI 2471/1-1 to U.D. as well as by the Research Center for Immunotherapy (FZI, Forschungszentrum für Immuntherapie) of the Johannes Gutenberg University Mainz.

## 6 Author Contributions

U.D. acquired and analyzed data, developed instrument methods, prepared figures. M.L. developed algorithms and the MIDIAID pipeline, analyzed data, prepared figures. M.S. developed algorithms and the MIDIAID pipeline, analyzed data, prepared figures. S.B. developed software, J.D. analyzed data, developed software. T.S. developed software.

J.K. analyzed data. F.K. developed instrument methods. O.R. analyzed data, developed instrument methods. D.T. and A.H. developed machine learning methods for edge refining. S.T. conceived the method, designed experiments, analyzed data. All authors discussed results and wrote the manuscript.

