## Supplementary Figures 1-4 for "midiaPASEF maximizes information content in data-independent acquisition proteomics"

### Supplementary Information

**V1.0 30.01.2023 biorXiv**

**Contents:**

Supplementary Figure 1:

Supplementary Figure 2

Supplementary Figure 3

Supplementary Figure 4

**Supplementary Figure 1:** Illustration of the total midiaPASEF isolation parallelogram (A) and zoom into the edge region (B) to visualize the overlapping window scheme for HeLa tryptic peptides. Data were analyzed using Bruker DataAnalysis software.

**A**

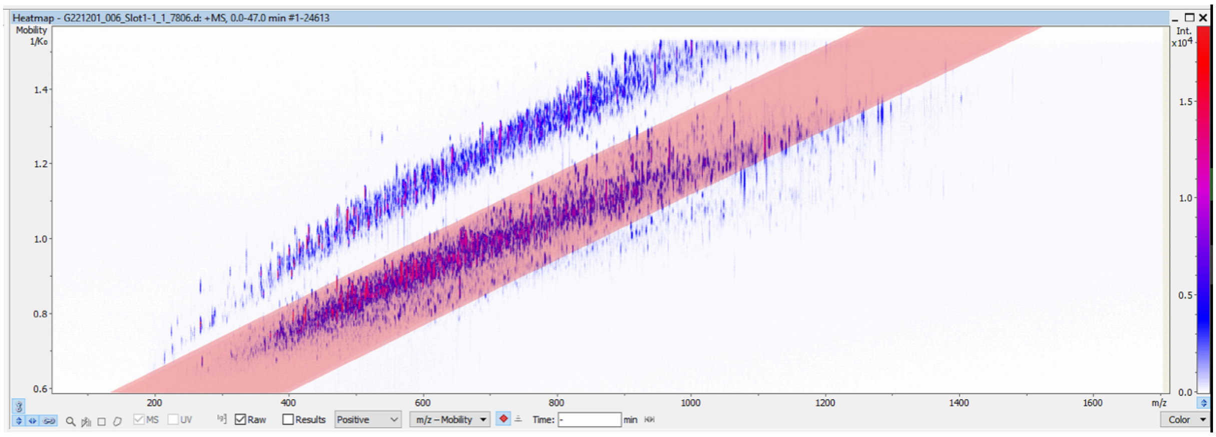


**B**

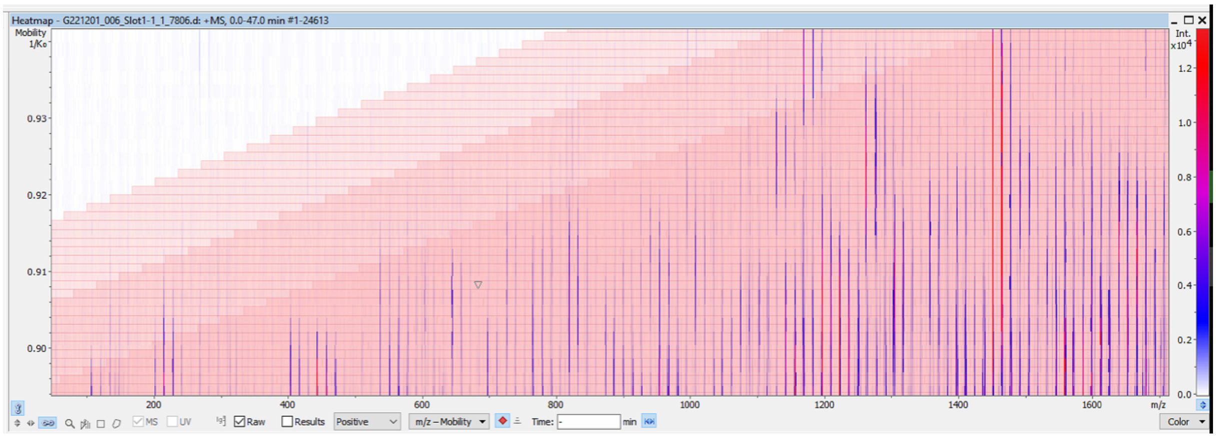

**Supplementary Figure 2:** Example annotated spectra of *de novo* Sequencing results of HeLa tryptic peptides using PEAKS X Pro v 10.6 . Rawdata were processed by the MIDIAID pipeline and resulting .mgf files *de novo* sequenced in PEAKS using 10 ppm precursor and 0.01 Da fragment ion mass tolerances.

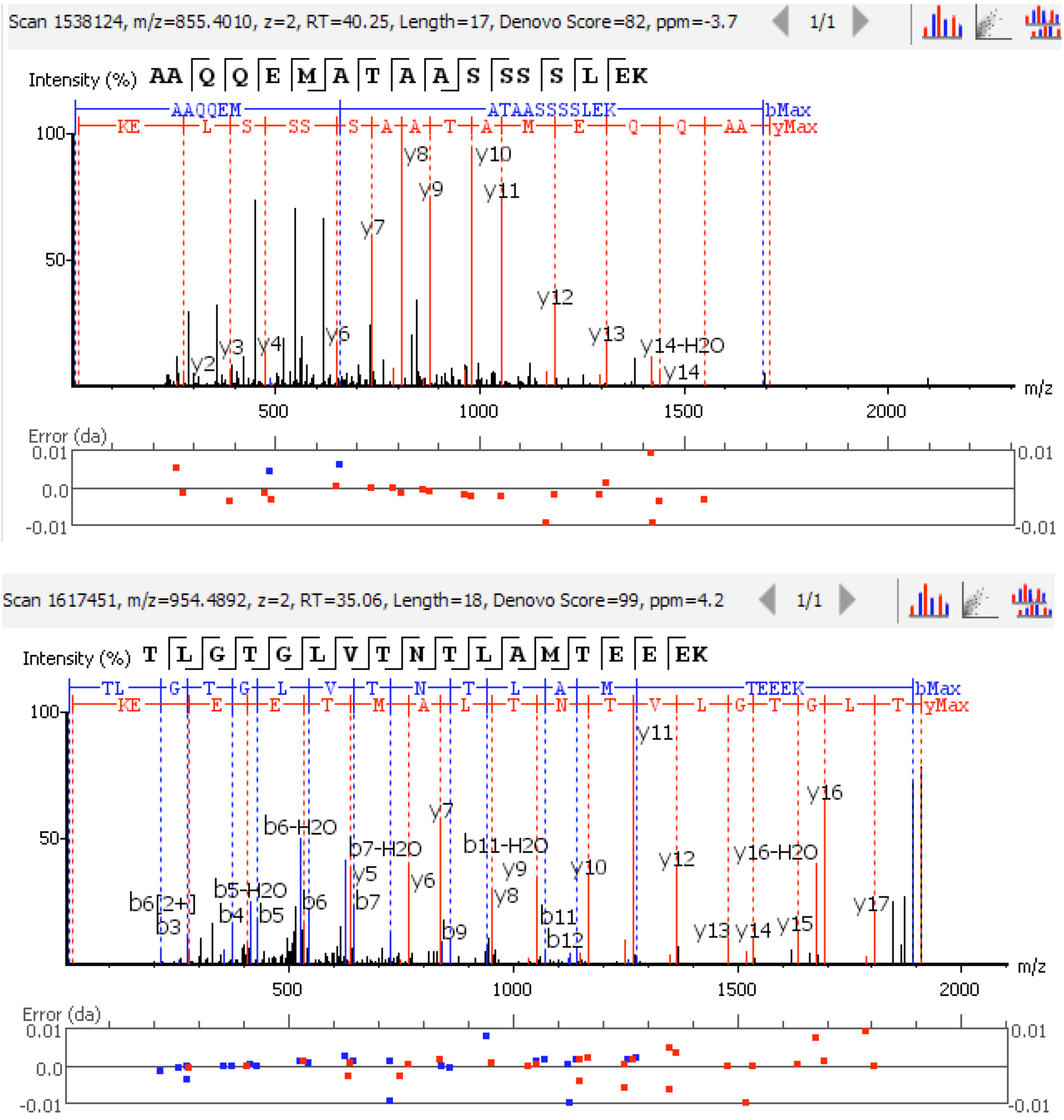

**Supplementary Figure 3**:

Overview of *de novo* sequencing results of deconvoluted midiaPASEF spectra obtained from HeLa tryptic peptides. Rawdata were processed by the MIDIAID pipeline and resulting .mgf files *de novo* sequenced in PEAKS using 10 ppm precursor and 0.01 Da fragment ion mass tolerances.

### 2. Result Statistics

**Figure 1.**

**(a)** Scatterplot of peptide Denovo score versus precursor mass error; **(b)** Distribution of residue local confidence in filtered result;

**(c)** Scatterplot of peptide ALC score versus ΔRT(difference of RT and predicted RT), **(d)** Scatterplot of peptide RT versus Predicted RT.

**(a)**

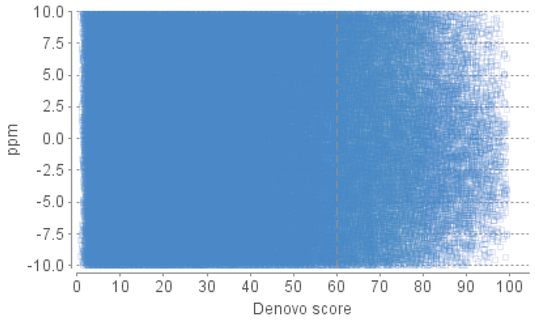


**(b)**

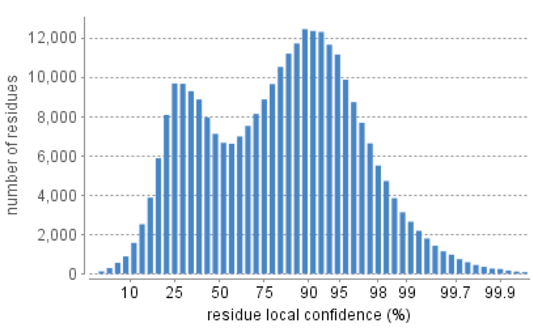


**(c)**

**No information available.**

**(d)**

**No information available.**

**Table 1.** Statistics of data and result.

| | |
|---|---|
| # of MS scans | 0 |
| # of MS/MS scans | 570721 |
| De novo peptides after score filter | 23899 |

**Table 2.** Result filtration parameters.

| | |
|---|---|
| De novo score | ≥60% |

**Supplementary Figure 4:** MIDIA enables library-free identification of MHC class II ligands. Example spectra of three MHC class II ligands identified from MHC class II immunoprecipitates from JY cells using 100 ms MIDIA acquisition. Rawdata were processed by the MIDIAID pipeline and resulting .mgf files search in PEAKS against human Uniprot database using 10 ppm precursor and 0.01 Da fragment ion mass tolerances.

A

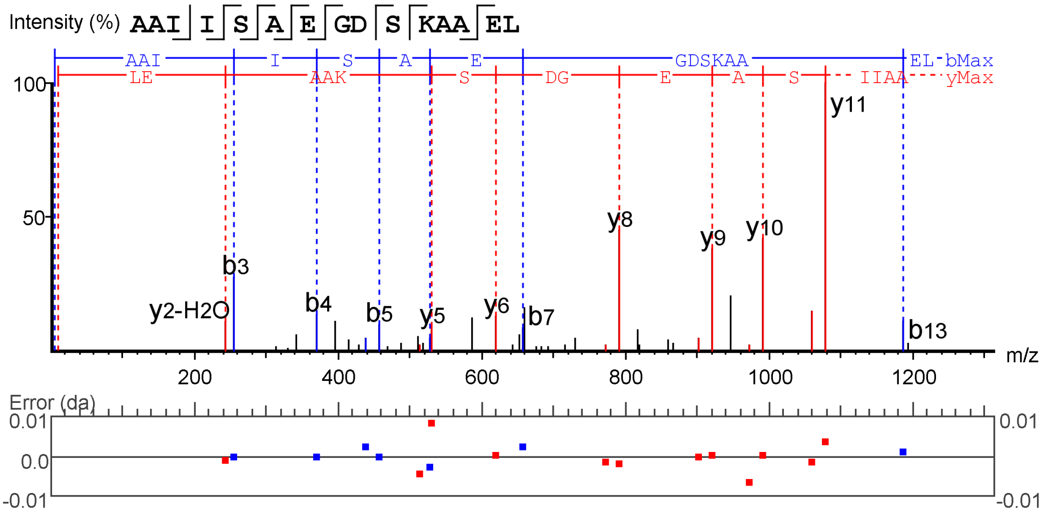


B

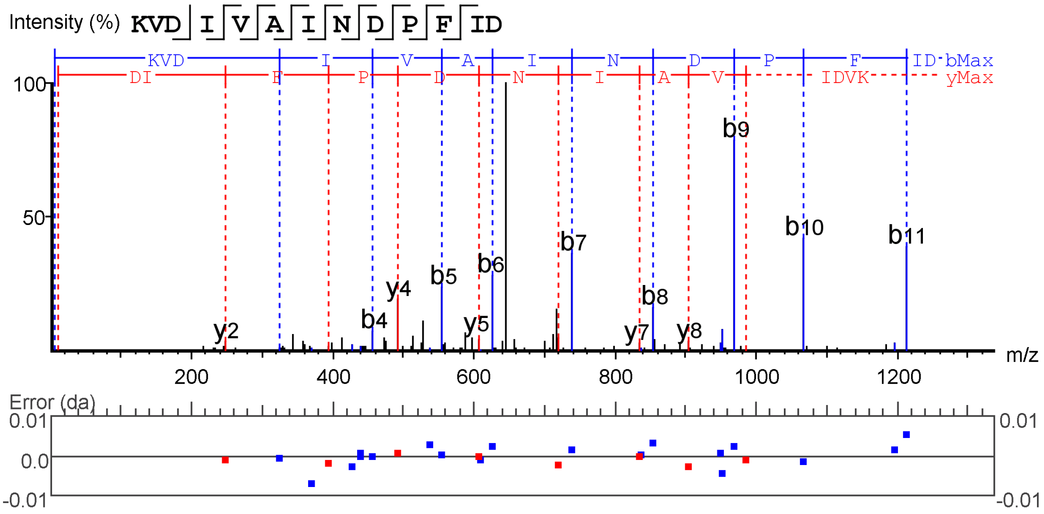


C

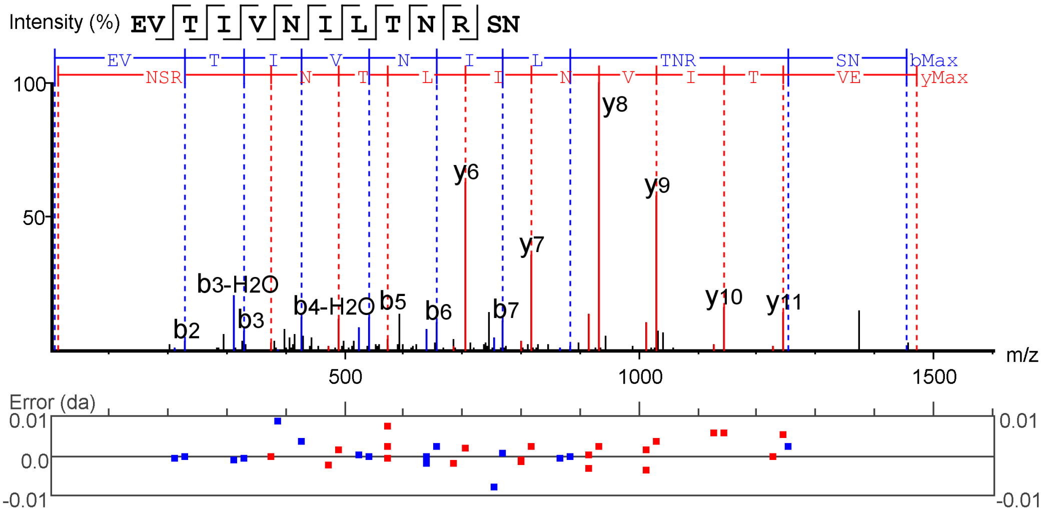
